# An aged human bone marrow mesenchymal niche causes impaired function of human hematopoietic stem cells under regeneration-induced stress

**DOI:** 10.64898/2026.10.27.762262

**Authors:** Ani Grigoryan, Ruba Abdelrahman, Wong Sik Seo, Alejandro Garcia-Garcia, Vadim Sakk, Antigone Karavelaki, Karin M Danzer, Christian Haupt, Hubert Schrezenmeier, Moritz Oltmanns, Heiko Reichel, Hartmut Geiger, Paul E Bourgine

## Abstract

Hematopoietic stem cells (HSCs) reside in the bone marrow (BM) and are responsible for blood formation throughout life. With advancing age, HSCs show reduced functional capacity. While clear intrinsic cellular changes in human HSCs (hHSCs) occur during aging, it remains unclear how the human bone marrow (hBM) niche contributes to hHSC dysfunction. This research gap is largely due to a lack of suitable experimental models capable of replicating the organization and function of the aged hBM environment. This study demonstrates that aged hBM mesenchymal stromal cells (hBMSCs) act as important niche regulators of hHSC function by contributing to the myeloid-biased differentiation of young hHSCs. Furthermore, hBMSCs play a crucial role in protecting hHSCs during regeneration-induced stress while this protective function of hBMSCs is impaired with age.

## INTRODUCTION

The hematopoietic system is maintained by HSCs located in BM niches (*1*). Within these niches, HSCs self-renew and differentiate to produce all blood cells (*1*). With age, hHSCs exhibit reduced self-renewal and reconstitution potential (*2-6*), which is clinically evident in BM transplantation, where HSCs from older donors result in poorer transplant outcomes associated with lower engraftment rates and higher overall mortality (*4, 5*). Moreover, aged hHSCs exhibit a tendency toward myeloid differentiation (*2, 6-8*), which is believed to contribute to the expansion of pre-malignant myeloid clones and, consequently, to the development of myeloid malignancies (*9, 10*). Given that many hematological disorders are age-related and that HSC aging is at least partially responsible for such conditions in the elderly, it is crucial to understand how hHSCs age and which factors drive stem cell aging in humans. This would open up opportunities to identify strategies to prevent, delay, or potentially even reverse hHSC aging.

Studies in mice have shown that the BM niche is a crucial regulator of HSC function (*11, 12*). It maintains stem cells and controls their differentiation (*11*). Also, mouse BM (mBM) niche changes with age, and these changes contribute to age-related HSC dysfunction (*13-17*). However, little is known about how aging affects the cellular and molecular composition of the hBM niche in humans and the contribution it makes to the aging of hHSCs (*18*). Although hematopoiesis is evolutionarily conserved between mice and humans, significant differences exist, for example regarding the formation of hematopoietic cell lineages (*19*). The surface markers used to isolate mouse HSCs (mHSCs) and hHSCs also differ, which may indicate functional differences in the mechanisms of stem cell-niche interaction that govern stem cell maintenance and differentiation (*19-22*). In xenotransplantation models, hHSCs exhibit a persistently high proliferation rate within mBM niches, in contrast to their predominantly quiescent state in hBM niches (*23, 24*). This suggests that the mBM niche differs in terms of the factors essential for the transition of hHSCs into a quiescent state, as the quiescent state is thought to be central for the proper regulation of HSC maintenance (*23, 24*). Therefore, novel models of hBM niches are required to investigate the function of hHSCs in vivo in the context of aging.

The ability of hBMSCs to establish a functional hBM niche, combined with advances in bone engineering strategies, has enabled the development of hBM niche models known as humanized ossicles (hOss) (*25-44*). Such hOss are formed in immunocompromised mice and have an architecture of a bone organ with mature cortical and trabecular bones and a well-developed BM cavity (*22, 26, 27, 36, 37, 45*). The hOss contain hBMSC populations in the form of osteoblasts, osteocytes, adipocytes and stromal cells, providing an hBM mesenchymal niche that offers superior support for healthy and malignant human hematopoiesis (*29, 36, 46*). However, the successful formation of hOss requires hBMSCs with excellent proliferation and chondrogenic differentiation capabilities - properties that are impaired with age (*47-49*). This age-related decline in the function of hBMSCs poses a major challenge for the creation of faithful models of the aged hBM mesenchymal niche (*26*). To overcome this limitation, this study presents a new strategy for generating aging-centered niche models from primary young and aged hBMSCs. This system provides the first in vivo platform modeling an aged hBM niche and its functional consequences on hHSCs.

## RESULTS

### Establishment of an ectopic hBM mesenchymal niche using young and aged hBMSCs

To bypass the age-associated performance variability of hBMSCs, a two-step protocol was established towards establishing an aged hBM niche model in vivo. First, a human mesenchymal cell line (MSOD-B cells) was primed to undergo chondrogenic differentiation in vitro to form a human hypertrophic cartilage tissue (**Fig.1A**), which was subsequently decellularized and freeze-dried to be used as a standardized and cell-free cartilage scaffold (**Fig.1A**) (*36, 44, 50*). Following subcutaneous implantation into immunocompromised NSG mice (**Fig.1B**), this cell-free human cartilage graft underwent rapid vascularization and remodeling and ultimately developed into an Oss within 4 weeks (**Fig.1B**) (*50-53*). In a second step, 500,000 primary hBMSCs from young (< 40 years, 5 donors) or older (> 65 years, 6 donors) donors were injected into this cell-free, cartilage-derived Oss to enable the formation of young or aged hBM mesenchymal niches, hereinafter referred to as young or aged hBM niches (**Fig.1C**). A gradual decline in hBM niche cells was observed (**Fig.1D**). By week 8, 50% of the Oss already exhibited a reduced number of human niche cells, and 16 weeks post-transplantation, the cells had disappeared from 50% of the Oss, while the remaining Oss contained only a few cells (**Fig.1D**). Nonetheless, both young and aged hBMSCs were able to engraft in mature Oss, and to undergo differentiation (**Fig.1E** and **fig.S1A-B**). This allowed the reconstitution of hBM niches with identified human bone-lining cells, osteoblasts, adipocytes, as well as stromal and blood vessel-associated cells (**Fig.1E** and **fig.S1A-B**), suggesting that the model enables the establishment of a mesenchymal niche regardless of donor age.

**Fig 1.**
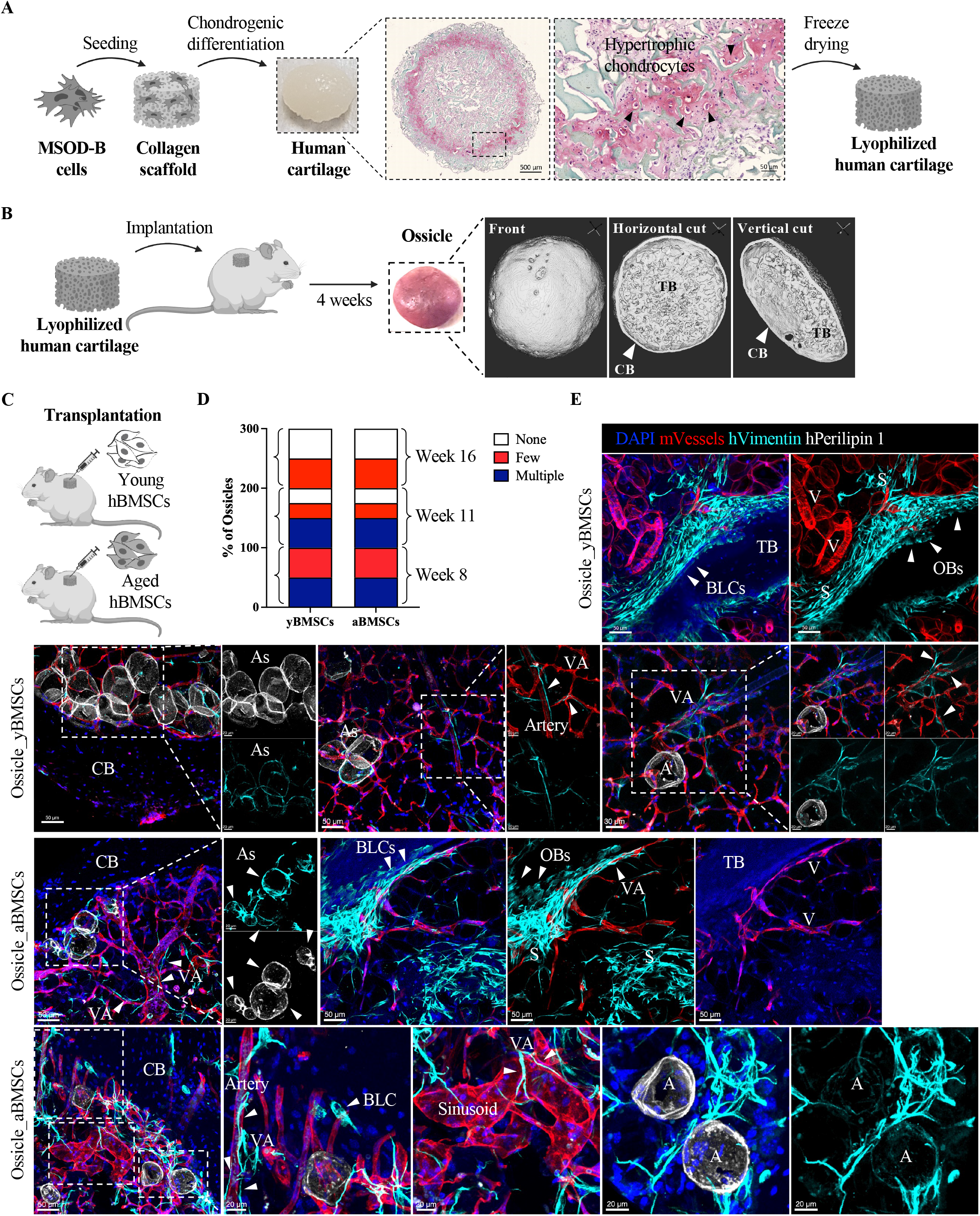
Establishment of an ectopic hBM mesenchymal niche using young and aged hBMSCs. (**A**) Schematic representation of the production of standardized, cell-free cartilage tissue; shown are the seeding and differentiation of a human mesenchymal cell line, a representative image of engineered human cartilage, Safranin-O staining of a human cartilage section, and the freeze-drying of the tissue. (**B**) Schematic representation of the subcutaneous implantation of these standardized and cell-free human cartilage tissues into immunocompromised NSG mice. A representative image from the micro-CT analysis of Oss formed after 4 weeks of in vivo implantation, showing mature cortical bone (CB) and trabecular bone (TB) as well as BM cavities. (**C**) Schematic representation of the transplantation (intraossicle injection) of primary hBMSCs from 5 young and 6 aged donors. (**D**) Confocal microscopy and immunostaining analysis of human mesenchymal cells in the Oss 8, 11, and 16 weeks after injection. “None” indicates Oss in which no human mesenchymal cells were detected; “few” refers to Oss exhibiting only scattered mesenchymal cells (predominantly stromal and vessel-associated cells) or cells in only localized tissue areas; “multiple” indicates Oss containing a large number of mesenchymal cells of various lineages, homogeneously distributed throughout the tissue sections. A total of 19 and 23 Oss, respectively, containing human mesenchymal cells from young and older donors, were analysed. (**E**) Representative images of human mesenchymal cells (young or aged) stained with a human-specific anti-vimentin antibody in 50-100 µm thick bone sections. Mouse blood vessels were stained with anti-laminin or anti-CD31 antibodies, while human adipocytes were stained with a human-specific anti-perilipin 1 antibody. Nuclei are stained with DAPI. Shown are cortical bone (CB), trabecular bone (TB), human bone-lining cells (BLCs), human osteoblasts (OBs), human adipocytes (As), and human vessel-associated (VA) and stromal cells.

### Young hHSCs exhibit functional changes in an aged hBM niche

To ensure the presence of hBM niche cells during hHSC transplantation and engraftment, hBMSCs and hematopoietic stem and progenitor cells (hHSPCs) were co-injected (**Fig.2A**). Human CD34^+^ HSPCs isolated from the BM of young, healthy donors were divided into two groups and mixed with either young hBMSCs or aged hBMSCs and injected into standardized, cell-free cartilage-derived Oss in sublethally irradiated NSG mice (**Fig.1A-B** and **Fig.2A**). A total of four young and four aged donors were used for the hHSPCs and the hBMSCs. 16 weeks after injection, the Oss were explanted for analyses. No differences regarding size, bone volume, or total number of BM cells were observed between the Oss injected with young hBMSCs (Oss_yBMSCs) and those injected with aged hBMSCs (Oss_aBMSCs) (**Fig.2B-E** and **fig.S2A**). However, there was a reduction in the number of BM mononuclear cells in Oss_aBMSCs (**Fig.2F**). Moreover, the overall hematopoietic engraftment rate was significantly lower in Oss_aBMSCs compared to Oss_yBMSCs (**Fig.2G-H**). Blood lineage reconstitution was also differed markedly, with young hHSCs in Oss_aBMSCs showing a reduced B cell and enhanced myeloid differentiation compared to those in Oss_yBMSCs (**Fig.2G** and **2I**). Importantly, this correlates with the myeloid tendency observed in peripheral blood (PB) analysis of young and older donors (**Fig.2J**). In addition, the rate of human hematopoietic engraftment in the mouse PB and in the femur carrying Oss-aBMSCs was lower, although no differences in differentiation were observed (**fig.S2B-G**). While no changes were detected in Lineage^-^CD38^-^CD34^+^ cells, as well as in multipotent (MPPs: Lineage^-^CD38^-^ CD34^+^CD90^-^CD45RA^-^) and lymphoid (LMPs: Lineage^-^CD38^-^CD34^+^CD90^-^CD45RA^+^) progenitor cells, the frequency of hHSCs (Lineage^-^CD38^-^CD34^+^CD90^+^CD45RA^-^) was significantly reduced in Oss_aBMSCs compared to Oss_yBMSCs (**Fig.2K-M** and **fig.S2H-I**). No changes were detected regarding hHSPCs in the femur (**fig.S2J-K**). Taken together, these data suggest that young hHSCs exhibit functional changes when exposed to an aged hBM niche (*2*).

**Fig 2.**
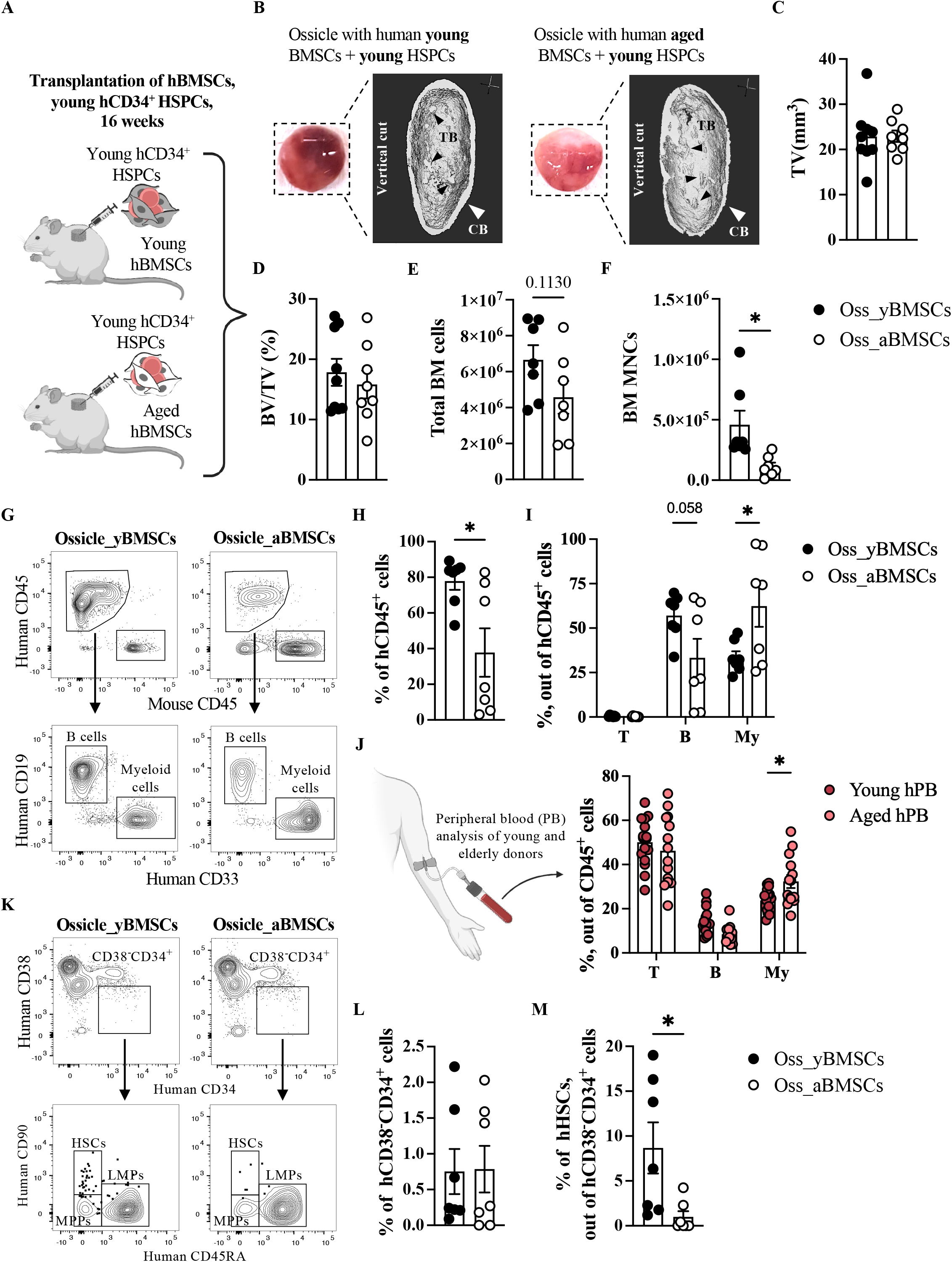
Young hHSCs exhibit functional changes in an aged hBM niche. **(A)** Schematic representation of intraossicle co-injection of hBMSCs (500.000 cells) and hHSPCs (250.000 cells) into Oss derived from standardized and cell-free cartilage grafts after 4 weeks of implantation. **(B)** Representative macroscopic images and micro-CT analysis images of Oss explanted 16 weeks after the injection of either young or aged hBMSCs together with young hHSPCs (referred to here as Oss_yBMSCs and Oss_aBMSCs, respectively). Cortical (CB) and trabecular (TB) bone as well as BM cavities are shown. **(C-D)** Quantification of total volume (TV) (C) and bone volume (BV; normalized to total volume) (D) of 9 Oss_yBMSCs and 8 Oss_aBMSCs. **(E-F)** Quantification of total BM (E) and mononuclear cells (F) of 7 Oss_yBMSCs and 7 Oss_aBMSCs. Statistical values were determined using *t* test; \**P* < 0.05. **(G)** Schematic representation of the flow cytometry gating strategy of human CD45^+^ (hCD45), mouse CD45^+^ (mCD45), human CD19^+^ B cells and CD33^+^ myeloid cells in 7 Oss_yBMSCs and 7 Oss_aBMSCs. **(H-J)** Flow cytometry analysis of hCD45^+^ chimerism (H) as well as human lymphoid (CD3^+^ T, CD19^+^ B) and human myeloid (CD33^+^ My) cells (I) in 7 Oss_yBMSCs and 7 Oss_aBMSCs, and in hPB from 16 young and 14 older donors (J). Statistical values were determined using *t* test and two-way ANOVA followed by Šidák’s test for multiple comparisons; \**P* < 0.05. **(K-M)** Schematic representation of the flow cytometry gating strategy (K) and analysis of hCD38^-^CD34^+^ cells within the hCD45^+^Lineage^-^ population (L), as well as hMPPs (CD90^-^CD45RA^-^), hLMPs (CD90^-^CD45RA^+^), and hHSCs (CD90^+^CD45RA^-^) (M) within the hCD38^-^CD34^+^ population in 7 Oss_yBMSCs and 7 Oss_aBMSCs. Statistical values were determined using *t* test; \**P* < 0.05. All analyses were performed 16 weeks after the intraossicle injection of hBMSCs and hHSPCs.

### A young hBM niche protects aged hHSCs from regeneration-induced stress

Conversely, to investigate the effects of young hBMSCs on the function of aged hHSCs, human aged CD34^+^ HSPCs were divided into two groups and mixed with either aged hBMSCs or young hBMSCs and injected into standardized and cell-free cartilage-derived Oss (**Fig.3A**). A total of 5 aged donors were used for the hHSPCs, and 3 young and 4 aged donors were used for the hBMSCs. Similar to the previous in vivo experiment, Oss were explanted 16 weeks post-transplantation for analysis (**Fig.3B**). No differences were observed regarding engraftment rate and differentiation (**Fig.3C-D**). However, the Lineage^−^CD38^−^CD34^+^ progenitor cell proportion was significantly higher in Oss_yBMSCs than in Oss_aBMSCs (**Fig.3E**), while LMPs and MPPs exhibited comparable frequencies (**Fig. 3F**). Most strikingly, aged hHSCs had completely disappeared in 5 of 6 Oss_aBMSCs, but were preserved in Oss_yBMSCs (**Fig. 3G**), similar to what was observed with young hHSCs (**Fig. 2M**). Taken together, these data suggest that a young hBM niche is able to maintain hHSCs during regeneration-induced stress, and that this ability is impaired with age.

**Fig 3.**
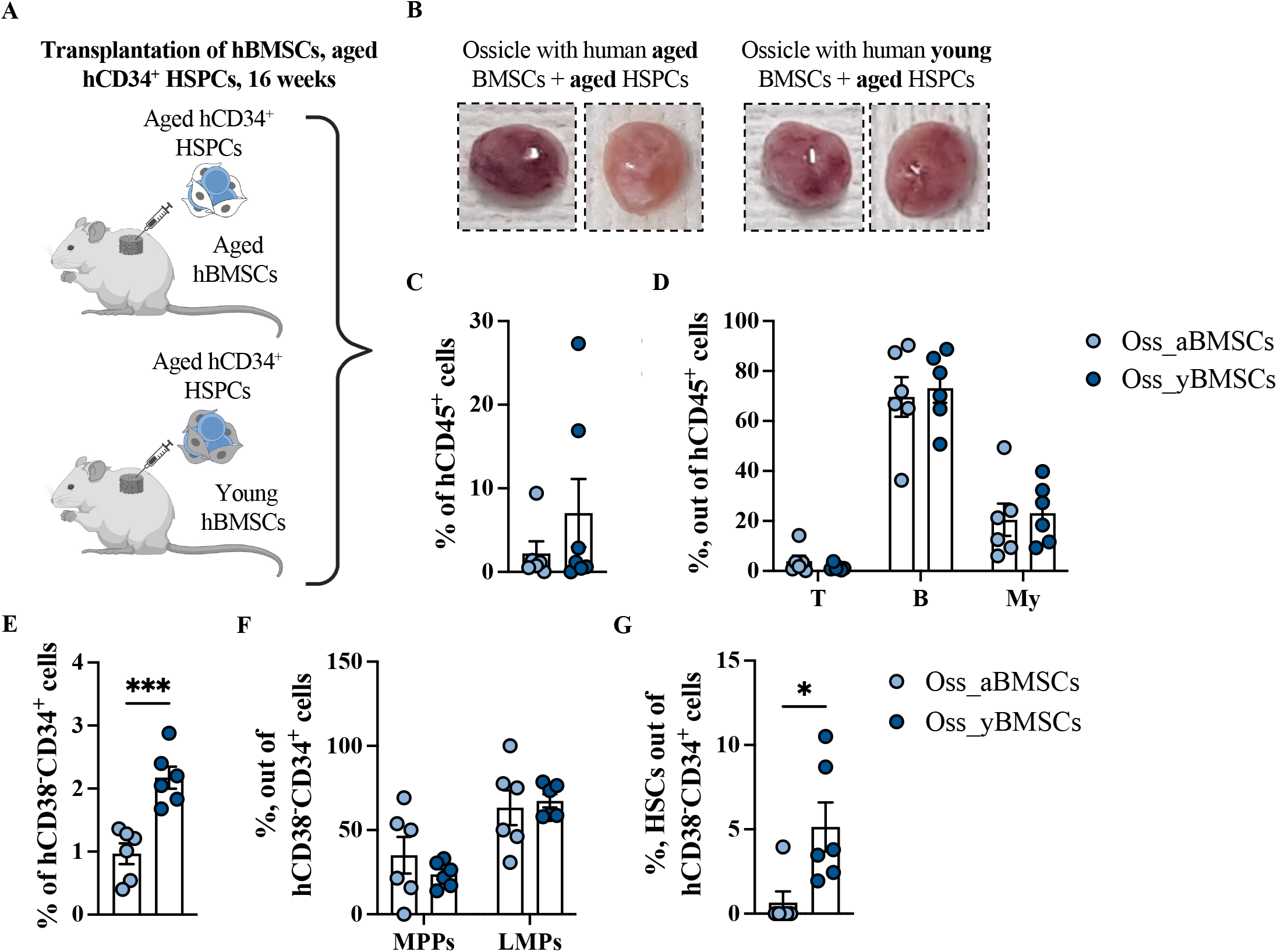
A young hBM niche protect aged hHSCs from regeneration-induced stress. (**A**) Schematic representation of intraossicle co-injection of hBMSCs (500.000 cells) and hHSPCs (58.000 cells) into Oss derived from standardized and cell-free cartilage grafts after 4 weeks of implantation. (**B**) Representative macroscopic images of Oss explanted 16 weeks after the injection of either aged or young hBMSCs together with aged hHSPCs (referred to here as Oss_aBMSCs and Oss_yBMSCs, respectively). (**C-D**) Flow cytometry analysis of hCD45^+^ chimerism (C) as well as human lymphoid (CD3^+^ T, CD19^+^ B) and human myeloid (CD33^+^ My) cells (D) in 6 Oss_yBMSCs and 6 Oss_aBMSCs. Statistical values were determined using *t* test and two-way ANOVA followed by Šidák’s test for multiple comparisons. (**E-G**) Flow cytometry analysis of hCD38^-^CD34^+^ cells within the hCD45^+^Lineage^-^ population (E), as well as hMPPs (CD90^-^CD45RA^-^), hLMPs (CD90^-^CD45RA^+^) (F), and hHSCs (CD90^+^CD45RA^-^) (G) within the hCD38^-^CD34^+^ population in 6 Oss_yBMSCs and 6 Oss_aBMSCs. Statistical values were determined using *t* test; \**P* < 0.05 or two-way ANOVA followed by Šidák’s test for multiple comparisons. All analyses were performed 16 weeks after the intraossicle injection of hBMSCs and hHSPCs.

## DISCUSSION

The BM niche, as a central regulator of HSC function, undergoes age-related changes that contribute to HSC dysfunction in aged animals (*13, 15, 16*). Uncovering these changes in humans has remained largely unresolved, due to the lack of relevant human experimental models. By exploiting a novel aging-focused hOss, our study demonstrates that an aged human mesenchymal compartment is sufficient to induce myeloid-biased differentiation of hHSCs in vivo. Importantly, this is in line with the clinically observed increase in myeloid-lineage cells in elderly donors (Fig. 2J), supporting the biological relevance of the age-associated phenotype observed in our experimental model. Our study also provides evidence that aged hBMSCs lose the ability to maintain both young and aged hHSCs under regeneration-induced stress. This finding has important clinical implications and may help explain the variability in the outcomes of HSC transplantation, which might be influenced not only by donor age but also by the fitness of the recipient’s BM niche (*4*). Our methodology is based on the establishment of Oss using standardized, cell-free human cartilage tissue and primary hBMSCs isolated from young or older donors (*50*). This provides the first in vivo platform for studying the functional consequences of the aging environment on hHSCs. However, a limitation of this model lies in its chimeric nature, as the vasculature and other BM elements are of mouse origin. Compared to native mouse bones, the distinctive feature of Oss lies in the formation of a complex human mesenchymal niche. In fact, the reconstituted hBM consists exclusively of mesenchymal elements, allowing us to specifically attribute observed age-related hHSC dysfunction to this human compartment. Nonetheless, the mesenchymal cell line(s) that significantly control the age-related changes in hHSCs still need to be identified. In addition, in our study cohort, age correlates with the anatomical harvest site and underlying cardiovascular or orthopedic conditions. Consequently, hBMSCs from healthy young donors were harvested from iliac crest aspirates, whereas aged hBMSCs originated from the sternum or femoral neck of older donors undergoing cardiac or hip replacement surgery. The hBMSCs harvested from femoral head and iliac crest have been shown to exhibit similar proliferation, differentiation and support of hBM CD34^+^ HSPCs in culture (*54*). The composition of hBM CD34^+^ HSPCs was also comparable in samples from the iliac crest and femoral head (*54*). Nevertheless, future in vivo studies using young and aged hBM samples from strictly identical anatomical sites would allow to formally disentangle chronological aging from putative anatomical location effects or comorbidities that influence hBMSC function.

## MATERIALS AND METHODS

### Study design

The overall goal of the study was to investigate the extent to which the hBM mesenchymal cells influence the function of hHSCs, using a model of human bone organs that recapitulates aging processes. To this end, we first generated standardized cartilage tissue in vitro from a human mesenchymal stromal cell line and implanted it into immunodeficient mice to facilitate the formation of bone organs. Subsequently, primary hBMSCs and hHSPCs from young or older donors were injected into these bone organs to reconstitute both the human mesenchymal niche and the human hematopoietic system. A period of 16 weeks post-transplantation was established as the experimental endpoint. No animals or bone organs were excluded from the study. The study was neither blinded nor randomized, and no statistical methods were used to determine sample size.

### Mice

NOD.Cg-*Prkdc*^*scid*^ *Il2rg*^*tm1Wjl*^/SzJ (NSG) mice were obtained from the Jackson Laboratories. All animal experiments at Lund University were conducted in accordance with the guidelines of the Lund University Animal Ethics Committee (15485-18, 19012-19). All animal experiments at Ulm University were performed in accordance with the German Animal Welfare Act (TierSchG) and the German Animal Welfare Laboratory Animal Regulations (TierSchVerV), which transpose EU Directive 2010/63/EU into national law. Approval for the animal experiments was reviewed and granted by the Tübingen Regional Council (Regierungspräsidium Tübingen) under reference number 1596. All animal housing and care at both universities were carried out in accordance with institutional guidelines, applying the 3R principles to minimize pain and distress.

### Human samples

Human bone marrow samples (aspirates and biopsies) were obtained from the iliac crest of 9 healthy young donors (aged 22-28 years, plus one 39-year-old) and from the sternum or femoral head-neck of 11 older donors (aged 65-81 years, plus one 53-year-old) undergoing cardiac surgery or hip replacement, respectively. Human peripheral blood samples were obtained from 16 young (aged 20-40 years) and 14 aged (aged 72-78 years) donors. The study was conducted in accordance with the Declaration of Helsinki. The study protocols were approved by the ethics committees of Lund University and Ulm University under approval numbers 2021-04046, 392/16, 357/23 and 66/24. Informed consent was obtained from all donors participating in the study.

### Cell isolation and expansion

The isolation of the total bone marrow cells - either from bone marrow aspirates or from bone marrow biopsies (by mincing, crushing, and filtering) - was performed using PBS at room temperature (RT). Density gradient centrifugation with Lymphoprep (STEMCELL Technologies) was performed to isolate BM mononuclear cells from the total BM cells (1600 rpm, 25 min, RT). Subsequently, CD34^+^ and CD34^−^ cells were isolated using a CD34 positive selection kit (Miltenyi Biotec) according to the manufacturer’s protocol, using cold PBS supplemented with 2% FBS and 2 mM EDTA (Sigma-Aldrich). After isolation, CD34^+^ cells were frozen in a mixture of 90% FBS and 10% DMSO for further use. To isolate BM-derived mesenchymal stromal cells (BMSCs), CD34^-^ cells were seeded at a density of ∼86,000 to 97,000 cells/cm^2^ in a complete medium; this consisted of α-Minimum Essential Medium (αMEM) supplemented with 10% fetal bovine serum (FBS), 1% penicillin-streptomycin-glutamine-ITS solution (100×), 1% HEPES (1 M), 1% sodium pyruvate (100 mM) (all from Gibco), and 5 ng/mL fibroblast growth factor 2 (FGF-2) (R&D Systems). Cultivation was carried out in an incubator at 37°C, 3% O_2_, and a humidified atmosphere. The medium was changed for the first time after 48 hours and subsequently twice weekly. Once the expanded stromal colonies had reached 80-90% confluence, the BMSCs were harvested as passage 0 cells using 0.05% trypsin-EDTA (Gibco) and frozen in a mixture of 90% FBS and 10% DMSO for further use. All experiments were performed using passage 1 BMSCs (expanded from passage 0 BMSCs, seeding density ∼5,700 cells/cm^2^, in a humidified incubator, at 37°C, 3% O_2_) and MSOD-B mesenchymal cell line (*50*) at passages 21-23 (seeding density ∼3200 cells/cm^2^, in a humidified incubator, at 37°C, 5% CO_2_) in complete medium.

### Generation of a standardized Oss

Following expansion, 3.5 million MSOD-B cells were seeded onto type I collagen scaffolds (cylinders 8 mm in diameter and 3 mm thick; Ultrafoam, Davol) - corresponding to a density of 3.5 × 10^3^ cells/cm^3^ - and cultured for 3 weeks in chondrogenic medium; this consisted of Dulbecco’s Modified Eagle Medium (DMEM) supplemented with 1% penicillin-streptomycin-glutamine, 1% HEPES (1 M), 1% sodium pyruvate (100 mM), 1% ITS (100×) (insulin, transferrin, selenium) (all from Gibco), 0.47 mg/mL linoleic acid, 0.12% bovine serum albumin (BSA), 0.1 mM ascorbic acid, 10^−7^ M dexamethasone (all from Sigma-Aldrich), and 10 ng/mL TGF-β3 (R&D Systems). The medium was changed twice weekly. After 3 weeks, MSOD-B-derived cartilage tissues were rinsed with PBS, transferred to 50 ml Falcon tube, snap-frozen in liquid nitrogen for 5 min, and subsequently lyophilized overnight at -80 °C and 0.05 mbar using Labconco FreeZone 4.5 Plus (Labconco) or Alpha2-4 LD Plus (Christ) freeze dryer. The lyophilized tissues were then stored at 4°C. Before subcutaneous implantation into 8-week-old NSG mice, lyophilized cartilage grafts were hydrated with PBS.

### Histological (Safranin O) staining

Human cartilage tissue derived from MSOD-B cells was washed in 1x PBS (Gibco) and fixed overnight at 4°C in 4% formaldehyde (PolyScience). The tissue samples were washed sequentially with 1x PBS and processed in an automated tissue processor (MICROM STP 120) according to the following program: a graded series of ethanol (Sigma-Aldrich) - twice at 70%, twice at 90%, twice at 99.9% - followed by three washes in xylene (Fisher Scientific) and finally paraffin (Surgipath Paraplast, Leica Biosystems). Each step lasted 1.5 hrs. Subsequently, the tissue samples were embedded in paraffin at an embedding station (MICROM 350-1 and 350-2) and sectioned at 10 µm using microtomes (MICROM HM 355S or Epredia HM 325). The sections were then dried overnight at RT. To remove the paraffin, the sections were washed twice for 7 min in xylene (Fisher Scientific) and then once for 3 min in a mixture of 99.5% ethanol and xylene (1:1 ratio). This was followed by rehydration of the sections in a graded series of ethanol (Sigma-Aldrich) at concentrations of 99.5%, 95%, 70%, and 35%, with each step performed twice for 7 min. The sections were stained for 3 min with Mayer’s hematoxylin solution (Sigma-Aldrich). A washing step in distilled water for 10 min followed to remove excess dye from the sections. Subsequently, staining was performed for 3 min using a 0.01% Fast Green solution (Thermo Scientific); excess dye was removed by briefly rinsing the sections with 1% acetic acid (Fisher Scientific) for 10 to 15 seconds. After staining the slides with a 0.1% Safranin O solution (Fisher Scientific) for 7-10 min, dehydration and clearing were carried out by immersing the slides in a series of ethanol solutions - 95% (4 min) and 99.5% (4 min) - followed by a mixture of 99.5% ethanol and xylene (1:1) (2 min). Finally, the sections were washed in xylene for 4 min to remove the ethanol and then mounted using the mounting medium Permount (Fisher Scientific).

### Microcomputed tomography (Micro-CT)

Micro-CT analysis of the Oss was performed using the U-CT system (MILabs, Netherlands) with a tungsten X-ray source operating at 50 kV and 0.21 mA. A full-rotation scan (360°) with an angular step of 0.250° was acquired for each sample. Volumes were reconstructed using MILabs analysis software with an isotropic voxel size of 10 µm. To analyze bone volume, the volume of highly mineralized tissue was quantified using Seg3D [v2.2.1, NIH, NCRR, Science Computing and Imaging Institute (SCI), University of Utah, USA]. For the total volume analysis, each sample was meshed using Blender (v2.82a, Netherlands) and analyzed via a custom-developed script (*44*).

### Immunofluorescence staining (IF)

The explanted Oss were fixed in 4% formaldehyde (PolyScience) at 4°C for 24 hrs and subsequently decalcified in 10% EDTA solution, pH = 8 (Sigma-Aldrich) at 4°C for up to two weeks with gentle agitation. Afterward, the Oss were embedded in cryo-embedding medium (Medite) and sectioned at a thickness of 50-100 µm using a Medite M360 cryostat. All staining and washing steps were performed with gentle agitation. The sections were blocked and permeabilized for 1 hr at RT using 0.5% Triton X-100 (Sigma-Aldrich) in PBS and 20% donkey serum (Jackson ImmunoResearch). Following blocking/permeabilization, staining was performed with primary antibodies overnight at 4°C and with highly cross-adsorbed secondary antibodies for 3 hrs at RT; between these steps, the samples were washed three times for 20 min with a solution of 0.1% Triton X-100 in PBS and 2% donkey serum. To eliminate unwanted fluorescence, the tissue sections were treated with the Vector TrueView Autofluorescence Quenching Kit (Vector Laboratories). The tissue sections were mounted using Vectashield mounting medium with DAPI (Vector Laboratories, H1200). Images of the samples were acquired using an LSM980 confocal microscope (Zeiss). Raw primary data were imported into the Imaris v10.1 software package for further processing and conversion into 3-dimensional images. The following primary antibodies were used: rabbit anti-laminin (Novus Biologicals, NB300-144), rabbit anti-human perilipin-1 (abcam, ab172907), mouse anti-human vimentin (abcam, ab8979), and goat anti-CD31 (R&D Systems, AF3628). The following secondary antibodies were used: AF488-donkey anti-mouse (Jackson ImmunoResearch, 715-545.150), AF594-donkey anti-rabbit (Jackson ImmunoResearch, 711-585-152), and AF647-donkey anti-goat (Jackson ImmunoResearch, 705-605-147).

### Cell transplantation and flow cytometry analysis

Following expansion, young and aged hBMSCs - either alone or mixed with hCD34^+^ HSPCs - were injected into the Oss of NSG mice. For hBMSC-hHSC transplantation, the mice received sublethal irradiation (200 cGy). PB chimerism was monitored via flow cytometry every 4 weeks for up to 16 weeks post-transplantation. After 16 weeks, the Oss and mouse femurs were harvested and BM cells were isolated by flushing the femurs with 21–22G needles or by crushing the Oss with micro-pestles and mincing them with scissors. Samples were collected in tubes containing PBS, 2% FBS, and 2 mM EDTA. Mononuclear cells were isolated via density gradient centrifugation using Lymphoprep (STEMCELL Technologies) and stained according to standard procedures. Sample analysis was performed on a LSR Fortessa (BD) or a Fortessa SORP flow cytometer (BD). The “Zombie UV Fixable Viability Kit” (BioLegend) was used to distinguish dead cells. Monoclonal antibodies against human CD45 (Alexa Fluor 700, clone HI30, BD, 560566) and mouse CD45 (FITC, clone 30-F11, BD, 553080) were used to differentiate between human and mouse cells. Human specific antibodies were used to analyze cell lineages in mouse and human PB and BM: CD3 (PE-Cy7, clone UCHT1, BD, 563423), CD19 (APC, clone HIB19, BD, 561742), and CD33 (PE, clone WM53, BD, 561816). Flow cytometric analysis results are presented as the percentage of CD19^+^ B cells, CD3^+^ T cells, and CD33^+^ myeloid (My) cells within the total population of donor-derived cells. For the analysis of early hematopoiesis, samples were stained with the following anti-human antibodies: CD38 (PE, clone HIT2, BD, 555460), CD34 (APC, clone 581, BD, 555824), CD90 (PE-Cy7, clone 5E10, BD, 561558), CD45RA (PerCP-Cy5.5 clone HI100, BD, 563429), and anti-human lineage cocktail (FITC, eBioscience, 22-7778-72). Flow cytometry data were presented as the percentage of hematopoietic stem cells (HSCs; gating: CD38^-^CD34^+^CD90^+^CD45RA^-^), multipotent progenitors (MPPs; gating: CD38^-^CD34^+^CD90^-^CD45RA^-^), and lymphoid-primed multipotent progenitors (LMPPs; gating: CD38^-^CD34^+^CD90^-^CD45RA^+^) within the donor-derived CD45^+^Lin^-^ cell population. Sample analysis was performed using FlowJo (version 10.10.1, BD).

### Statistical analysis

Data are presented as mean ± SEM unless otherwise stated. Statistical analyses were performed using GraphPad Prism v11.1.0. Each experiment was conducted with at least three biological replicates; exact numbers are provided in the respective figure legends. The *t* test was used for comparisons between two groups, whereas analysis of variance (ANOVA) combined with a post-hoc test for multiple comparisons was used for comparisons between more than two groups and for multifactorial analyses. Statistical tests and *P* values are indicated in the respective figure legends. *P* values < 0.05 were considered significant.

## Acknowledgments

We thank the FACS and imaging facilities of the Lund Stem Cell Center, as well as the Lund University Bioimaging Center, for the X-ray CT system. We thank the Animal Research Centre of Ulm University for their support in handling and housing the mice. We also thank Prof. Dr. Stefan Scheding (Division of Molecular Hematology, Lund University) for providing bone marrow aspirates from young donors. The experimental setups were visualized using BioRender.com.

## Author contributions

Conceptualization: A.G., P.E.B.

Methodology: A.G., R.A., W.S.S, A.G.G., V.S., A.K.

Resources: A.G., P.E.B., K.M.D., C.H., H.S, M.O., H.R., H.G.

Investigation: R.A.,

W.S.S., A.G.G., A.K., A.G.

Visualization: A.G.

Supervision: P.E.B., A.G.

Writing – original draft:

A.G. Writing - review and editing: all authors.

## SUPPLEMENTARY MATERIALS

Figs. S1 to S2

**Fig S1.**
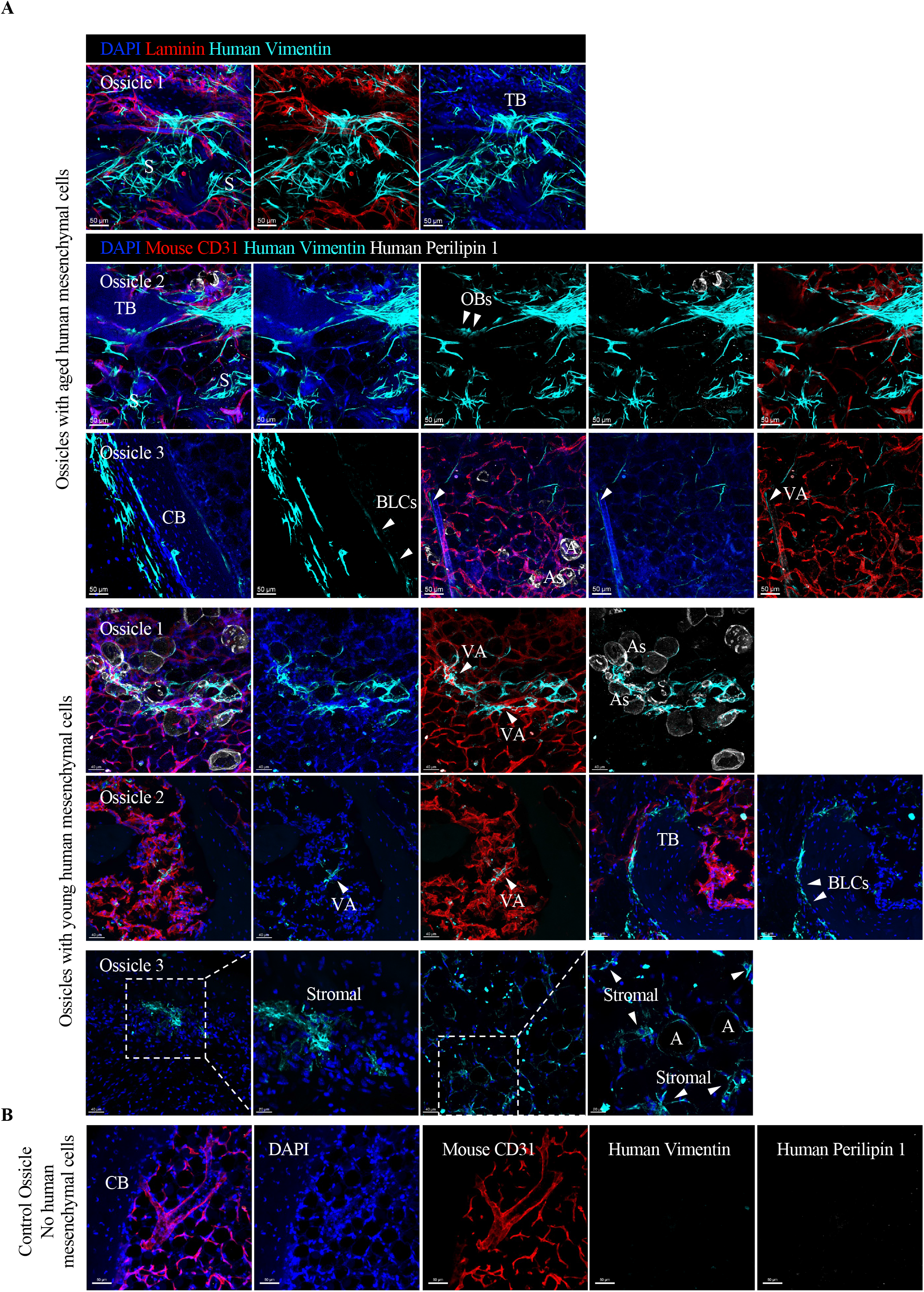
Establishment of an ectopic hBM mesenchymal niche using young and aged hBMSCs. (**A**) Representative images of human mesenchymal cells (young or aged) stained with a human-specific anti-vimentin antibody in 50-100 µm thick bone sections. Mouse blood vessels were stained with anti-laminin or anti-CD31 antibodies, while human adipocytes were stained with a human-specific anti-perilipin 1 antibody. Nuclei are stained with DAPI. Shown are cortical bone (CB), trabecular bone (TB), human bone-lining cells (BLCs), human osteoblasts (OBs), human adipocytes (As), and human vessel-associated (VA) and stromal cells. (**B**) Representative image of a 50-100 µm thick bone section from a control Oss not injected with hBMSCs, stained with anti-mouse CD31, anti-human vimentin, and anti-human perilipin-1 antibodies, demonstrating the specificity of the human antibodies.

**Fig S2.**
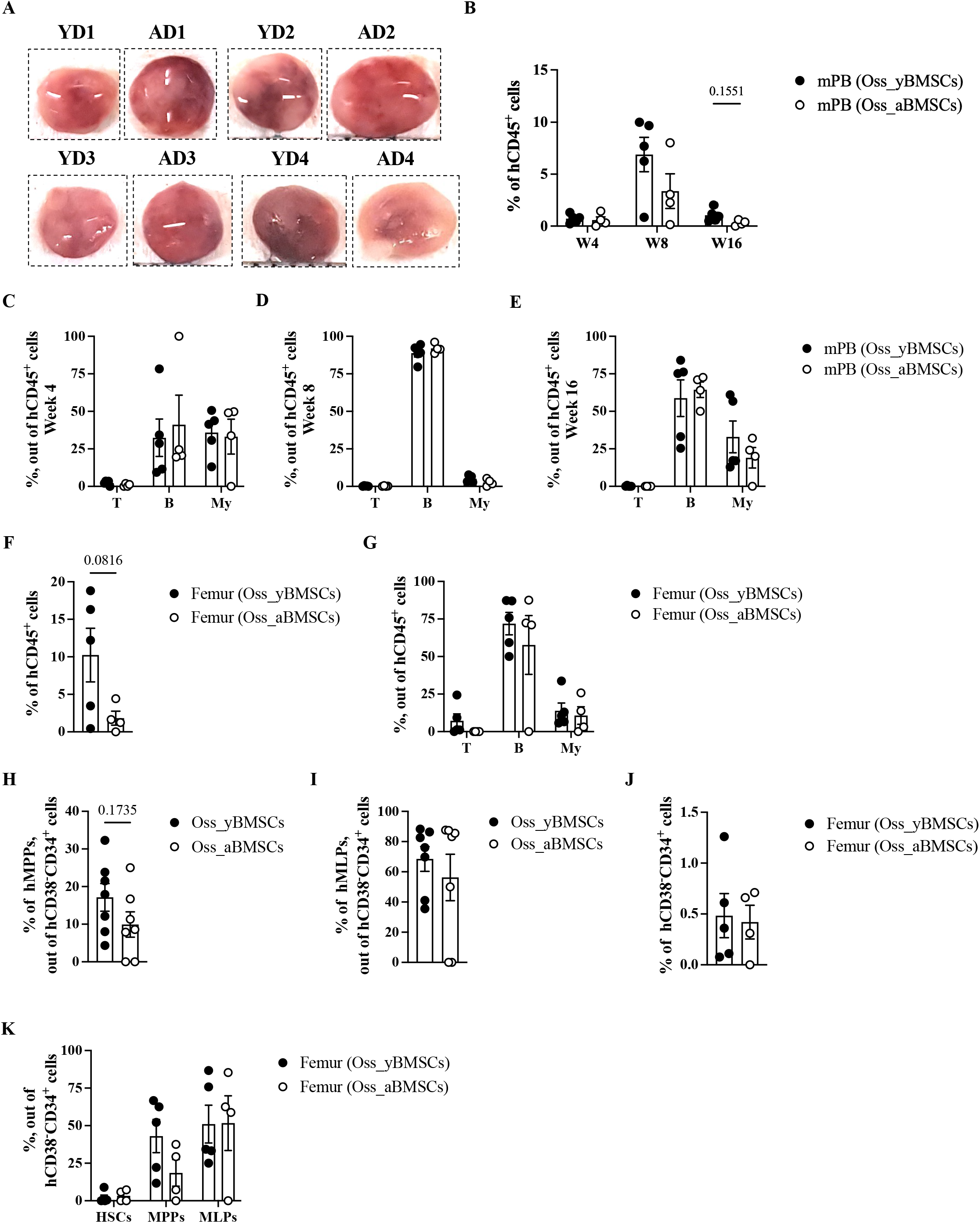
Young hHSCs exhibit functional changes in an aged hBM niche. **(A)** Representative macroscopic images of Oss explanted 16 weeks after the intraossicle injection of hBMSCs and hHSCs. **(B-E)** Flow cytometry analysis of hCD45^+^ chimerism (B) as well as human lymphoid (CD3^+^ T, CD19^+^ B) and human myeloid (CD33^+^ My) (C-E) cells in mouse PB (mPB) 4 (C), 8 (D) and 16 (E) weeks after transplantation, n=4-5 mice. Statistical values were determined using *t* test or two-way ANOVA followed by Šidák’s test for multiple comparisons. **(F-G)** Flow cytometry analysis of hCD45^+^ chimerism (F) as well as human lymphoid (CD3^+^ T, CD19^+^ B) and human myeloid (CD33^+^ My) (G) cells in the mouse femur 16 weeks after transplantation. Statistical values were determined using *t* test or two-way ANOVA followed by Šidák’s test for multiple comparisons. **(H-I)** Flow cytometry analysis of hMPPs (CD90^-^CD45RA^-^) (H) and hLMPs (CD90^-^ CD45RA^+^) (I) within the hCD38^-^CD34^+^ population in 7 Oss_yBMSCs and 7 Oss_aBMSCs. Statistical values were determined using *t* test. **(J-K)** Flow cytometry analysis of hCD38^-^CD34^+^ cells (J) within the hCD45^+^Lineage^-^ population, as well as hMPPs (CD90^-^CD45RA^-^), hLMPs (CD90^-^CD45RA^+^), and hHSCs (CD90^+^CD45RA^-^) (K) within the hCD38^-^CD34^+^ population in mouse femur 16 weeks after transplantation, n=4-5 mice. Statistical values were determined using *t* test or two-way ANOVA followed by Šidák’s test for multiple comparisons.

